# Downregulation of E-cadherin in pluripotent stem cells triggers partial EMT

**DOI:** 10.1101/2020.05.18.101899

**Authors:** C. Aban, A. Lombardi, G. Neiman, M.C. Biani, A. La Greca, A. Waisman, L.N. Moro, G. Sevlever, S. Miriuka, C. Luzzani

**Affiliations:** LIAN-CONICET, FLENI. Ruta 9, Km 53, Belén de Escobar (B1625), Buenos Aires, Argentina

**Keywords:** E-cadherin, Pluripotent Stem Cells, Partial EMT

## Abstract

Epithelial to mesenchymal transition (EMT) is a critical cellular process that has been well characterized during embryonic development and cancer metastasis and it also is implicated in several physiological and pathological events including embryonic stem cell differentiation. During early stages of differentiation, human embryonic stem cells pass through EMT where deeper morphological, molecular and biochemical changes occur. Though initially considered as a decision between two states, EMT process is now regarded as a fluid transition where cells exist on a spectrum of intermediate states. In this work, using a CRISPR interference system in human embryonic stem cells, we describe a molecular characterization of the effects of downregulation of E-cadherin, one of the main initiation events of EMT, as a unique start signal. Our results suggest that the decrease and delocalization of E-cadherin causes an incomplete EMT where cells retain their undifferentiated state while expressing several characteristics of a mesenchymal-like phenotype. Namely, we found that E-cadherin downregulation induces *SNAI1* and *SNAI2* upregulation, promotes *MALAT1* and *LINC-ROR* downregulation, modulates the expression of tight junction occludin 1 and gap junction connexin 43, increases human embryonic stem cells migratory capacity and delocalize b-catenin. Altogether, we believe our results provide a useful tool to model the molecular events of an unstable intermediate state and further identify multiple layers of molecular changes that occur during partial EMT.

## Introduction

Human pluripotent stem cells have the unique capacity to differentiate into the three germinal layers of the embryo (mesoderm, endoderm and ectoderm) and can build up all cell lineages of the body. In fact, formation of the trilaminar embryonic disk requires sequential rounds of differentiation of these cells into different and specialized cell types (1) demonstrating their high versatility. In particular, human embryonic stem cells (hESC) are pluripotent stem cells (PSC) derived from the inner cell mass of blastocyst and are able to differentiate into diverse cellular types using known stimuli. Nowadays, differentiation of mesenchymal stem cells from PSC represents a valuable strategy due to their relevant immunomodulatory properties and offers a renewable source of these cells. For this, several protocols and strategies have been established during the past years (2, 3).

PSC remain closely associated with neighboring stem cells growing in compact colonies. However, during early stages of differentiation, PSC undergo through an epithelial to mesenchymal transition (EMT) where deep morphological, molecular and biochemical changes occur (4, 5). EMT involves a series of events where epithelial cells lose their adhesion properties and acquire migratory capacity and other traits of a mesenchymal phenotype (1). This biological process has been comprehensively characterized and its implicated in several physiological and pathological events including embryonic stem cell differentiation, tissue repair and acquisition of certain properties of cancer stem cells (6).

Hallmark changes of EMT include alterations in cytoskeleton architecture, acquisition of migratory capacity, loss of apico-basal polarity and loss of cell adhesion. This phenotypic switch is mediated by activation of master transcription factors of EMT including SNAI1, SNAI2 and ZEB1/2 whose functions are finely regulated at transcriptional and translational levels (7). In line with this, initiation of EMT involves changes in gene expression and activation of signaling pathways (8). Also microRNAs, long noncoding RNAs and modification at post-translational levels are implicated in the regulation of this process (9). However, one of the main initiation signals of EMT is downregulation of E-cadherin (CDH1), a homophilic protein that mediates attachment to neighboring cells through interaction of its extracellular domains. Its expression is decreased during EMT and also loss of function of this protein promotes this transition. The transcriptional repression of E-cadherin has long been considered a critical step during EMT (10).

Initially, EMT was considered as a decision between either epithelial or mesenchymal states. Nowadays, it is well known that EMT is not a binary process and that it can be considered as a fluid transition where cells exist on a spectrum of intermediate states. During EMT, cells can adopt a hybrid and transient state called partial EMT phenotype (9). Cells in partial EMT, also known as an incomplete EMT, have characteristics of both epithelial and mesenchymal phenotypes. These mixed traits enable cells to undergo collective cell migration instead of an individual cell migration occurring in mesenchymal cells (11, 12). Partial EMT has been usually labeled as a metastable and reversible state (12, 13), which is determined by cell type and signals strength that initiate and maintain continuity of EMT process.

In this work we describe the molecular effects of E-cadherin downregulation in human embryonic stem cells, as a unique signal for inducing EMT. Our results suggest that induced decrease of E-cadherin expression by CRISPR interference system lead to partial EMT obtaining cells that conserve their pluripotency but also express characteristics of a mesenchymal-like phenotype.

## Methods

### Embryonic stem cells culture

Human embryonic stem cells HES3 were generously donated by Edouard Stanley’s Lab. Cells were maintained in an undifferentiated state under feeder-free conditions on GelTrex-coated plates (diluted 1:1000 from 15 mg/ml) (Thermo Fisher Scientific, USA). Cells were fed daily with mTeSR medium (STEM CELL Technologies, Canada) and maintained at 37°C in a humidified atmosphere with 5% CO2. When cells reached approximately 70-80% of confluence, colonies were dissociated using TrypLE 1x (Thermo Fisher Scientific, USA) and cells were seeded on new culture dishes previously coated with Geltrex in mTeSR medium (STEMCELL Technologies, Canada) containing 5 μM ROCK Inhibitor (Y-27632). For experiments, cells were seeded in mTeSR medium. Once cells attached to the plate (typically after 2 h), medium was changed and doxycycline (Dox) (Sigma Aldrich, USA) was added. Cells were incubated with Dox during 48, 72 and 96 h and Dox was renewed daily with medium change. At each of the indicated time points, material was collected to be analyzed later.

### Generation of stable cell line

For inducible CDH1 (E-cadherin gene) downregulation, a stable cell line called HES3-KRAB^CDH1^ was generated using the CRISPR interference system. To achieve this, a sgRNA directed to the promotor (200 bp upstream of TSS) of CDH1 was cloned into the pLenti-Sp-BsmBI-sgRNA-Puro vector (Addgene plasmid number 62207) and used to generate non-replicative lentiviral particles, as previously described by Neiman et al (2019). Sequences of sgRNA are listed in supplemental material (Table S1). Briefly, HEK293FT cells were co-transfected with sgRNA and lentiviral packaging plasmids and viral supernatants were collected at 48 h. Parental HES3 cells that express TRE-dCas9-KRAB construct (Neiman et al 2019) were dissociated to single cells with TrypLE and seeded at low density in mTser medium on GelTrex coated 6-well plates. On the following day, viral supernatant was added to the culture medium for 48 h. Then, medium was replenished and cells were treated with 1 μg/ml puromycin for 2 days. After selection, clonal isolation was performed. For this, 1 μl of a highly diluted cell suspension was plated in a 6-well plate and single cells were incubated with ROCK inhibitor (Y-26732) until they could form small colonies. After 7 days, individual colonies were hand-picked into 24 well plates and treated with 500 ng/ml of Dox to induce KRAB expression and E-cadherin repression. Cells were PCR genotyped for dCas9-KRAB and E-cadherin. Primer sequences are detailed in Table S2 (supplementary material).

### Immunofluorescence staining

Cells were seeded on coverslips coated with GelTrex in 24 well-plate and treated with Dox at different times. Then cells were fixed with 4% paraformaldehyde for 30 minutes, washed three times with PBS (Sigma Aldrich, USA) and blocked for 45 minutes with 0,1% Triton X-100 (Sigma Aldrich, USA), 0,1% bovine serum albumin (BSA, Gibco, USA) and 10% Normal Goat Serum (Gibco, USA) in PBS at room temperature. Then, cells were washed twice with PBS for 5 minutes and incubated with specific primary antibodies overnight at 4°C. The following day, cells were washed three times for 5 minutes and incubation with Alexa-fluor-conjugated secondary antibodies (Alexa 488 and Alexa 594, Life Technologies, USA) was performed for 1 hour in a dark humid chamber. After that, cells were washed three times for 5 minutes with PBST (PBS + 0,1% Triton X-100). Nucleus were counterstained with DAPI for 15 minutes. Finally, cells were mounted on glass slides and examined by fluorescence microscopy on an EVOS Digital Color Fluorescence Microscope (Thermo Fisher Scientific, USA). Primary antibodies used in this work include anti- CDH1 (1:400, Clone 36/E-cadherin; Cat 612131, BD Biosciences, USA), anti- HA-tag for KRAB protein (1:500, 32-6700 NOVEX HA epitope Tag antibody 5B1D10, Invitrogen, USA) and anti-b-Catenin (1:400, 4D5, MA5-15569, Invitrogen, USA).

### Quantitative qPCR analysis

Total RNA was isolated using Trizol Reagent (Invitrogen) according to manufacturer instructions and reverse-transcribed using M-MLV Reverse Transcriptase (Promega, USA). cDNA was amplified by qPCR with FastStart Universal SYBR Green Master reaction mix (Roche, Thermo Fisher Scientific, IN, USA) using specific primers detailed in Table S2 (supplementary material). qPCR was performed using a StepOnePlus Real Time PCR System (Applied Biosystems, USA) and analyzed using LinReg Software. All gene expression results were normalized to the geometric mean of RPL7 and HPRT1 housekeeping genes for each condition. Gene expression was analyzed by technical duplicates using at least three independent biological replicates.

### Wound healing assay

Cells were seeded at low confluence in a 24 well-plate. After different days of dox treatment, when cells reach 100% confluence, scratch wound healing assay was performed. Confluent monolayer was scratched with a sterile p200 pipette tip and immediately, cells were washed twice with PBS and fresh medium with Dox was added. Images were acquired at 0 and 22 h after injury via EVOS microscope (Life Technologies, USA). Experiments were performed at least three times. Quantification of the covered area was done by ImageJ software.

### Quantification of cell area

Bright field images were acquired with EVOS microscopy (Thermo Fisher Scientific, USA). Images of three independent experiments were evaluated for each condition (control without Dox and Dox treatment at 96 hours). The Cellpose segmentation algorithm (14) was use to create a mask where each cell is individually delineated. This mask was then used to quantify the area of each cell using the ImageJ-FIJI software.

### Statistical analysis

Results were expressed as mean ± S.D. of at least three biological replicates. Statistical significance was determined using t-test and ANOVA. Comparison of means between groups was assessed using Tukey test. Residual fitted normal distribution and homogeneity of variance. Statistical differences were referred to control conditions. Results were considered significant when p<0.05.

## Results

### Characterization of HES3-KRAB^CDH1^ cells

EMT is a key biological process that occurs during embryogenesis, normal wound healing and cancer metastasis. Accordingly, a plethora of stimuli can trigger this event and most of them converge in a downregulation of E-cadherin expression, a key process of EMT. To study whether downregulation of E-cadherin is sufficient to induce an EMT-like process we first generated a hESC line that represses this gene using a catalytically inactive CRISPR/Cas9 fused to a KRAB repressor domain inducible by Dox and a sgRNA targeting the *CDH1* promoter that encodes E-cadherin (15)(Fig. 1A). After establishing the clonal cell line named HES3-KRAB^CDH1^, we validated its effectiveness. Expression of KRAB protein was evaluated by immunofluorescence after cells were incubated with Dox (500 nM) during 48, 72 and 96 h. Without Dox, cells did not express KRAB protein but after Dox addition, positive immunostaining of KRAB is detectable and increased gradually during incubation time (Fig. S1). Then, we analyzed the efficiency of transcriptional repression of this cell line. For this, expression level of E-cadherin gene (*CDH1*) was measured by qPCR after cells were incubated with Dox at different times. As soon as 48h, E-cadherin mRNA levels decreased significantly but a further reduction was observed at 72 and 96h, times at which E-cadherin levels showed a 3-fold reduction relative to control condition (Fig. 1B). Parental HES3 wild type cells displayed no changes in expression of this gene after Dox treatment (Fig. S2). Finally, we assessed the inducible repression of E-cadherin at protein level. In the absence of Dox, as expected, E-cadherin was mainly localized at cell membrane. Upon Dox treatment, E-cadherin levels substantially decreased in a time-dependent manner, until its levels were almost undetectable at 96 h (Fig 1C). Overall, these results demonstrate that this inducible hESC line efficiently downregulates E-cadherin both at transcriptional and protein level.

**Fig. 1.**
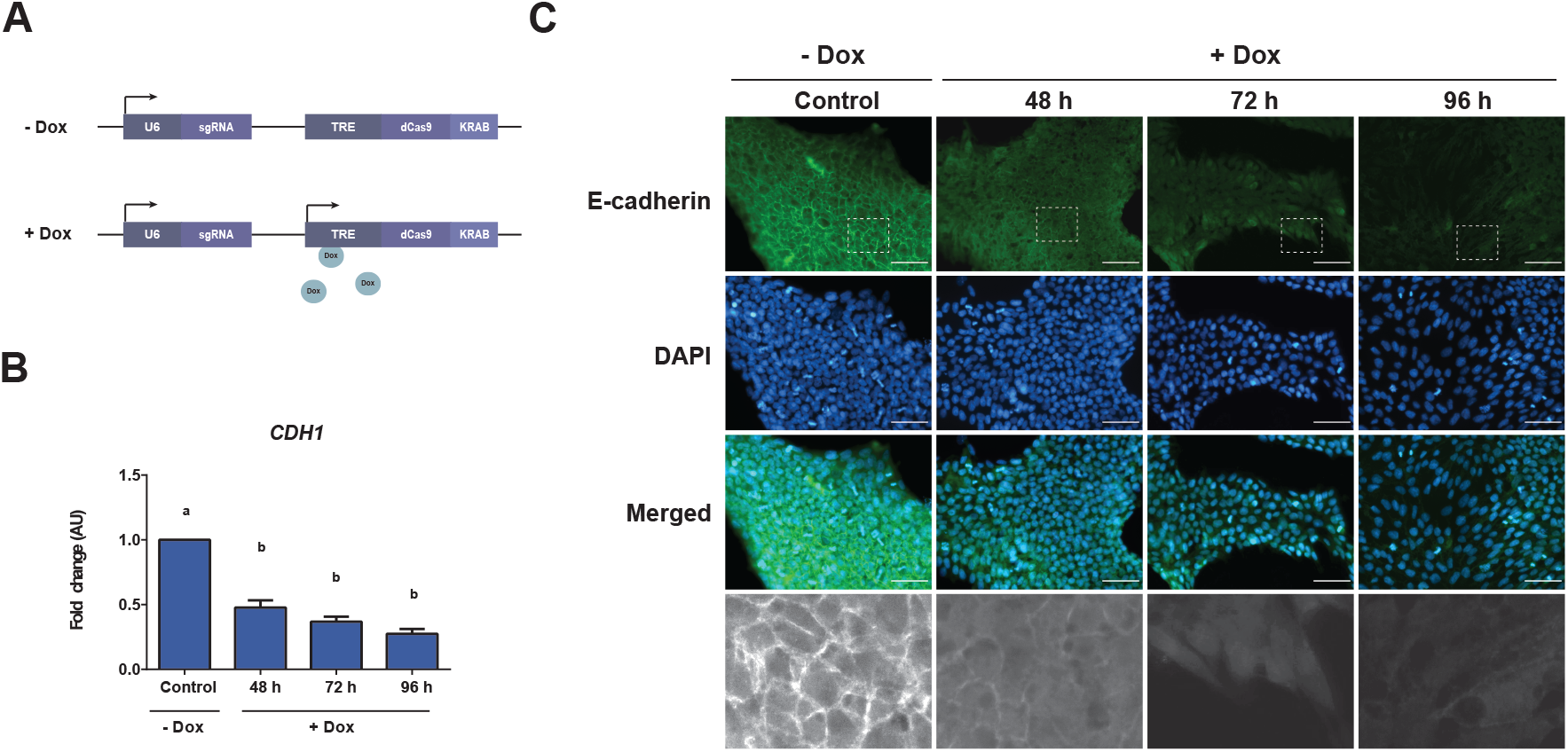
Characterization of HES3-KRAB ^CDH1^ clonal cell line. A) Schematic representation showing the construct used for generation of HES3-KRABCDH1 cell line. This knockdown system contains a deactivated Cas9 (dCas9) fused to a transcriptional repressor domain Krüppel Associated Box (KRAB) under the control of TRE promoter inducible by Dox. A sgRNA against CDH1 promoter was placed under the control of the U6 promoter which is constitutively activated. In cells not treated with Dox, the system is not active. Upon Dox induction, dCas9-KRAB is expressed and targeted to CDH1 promoter by the sgRNA to produce transcriptional repression of E-cadherin. B) Relative mRNA levels of E-cadherin (CDH1) in cells after incubation with Dox at different times assessed by qPCR. AU is the abbreviation for arbitrary units. Results are represented as mean ± SD (n=5). Different letters indicate significant differences of groups compared to control condition (p<0,001) for ANOVA with post hoc Tukey. C) Immunofluorescence images of E-cadherin protein (green) in cells incubated with Dox during 48, 72 and 96 h. Nuclei were stained with DAPI (blue). Magnification of the indicated areas are shown in black and white at the bottom of the panel. Representative images of at least three experiments are shown. Scale bar 50 μm.

### Analysis of intercellular junctions after E-cadherin downregulation

Considering the structural relevance of E-cadherin in cell structure, we next assessed whether changes in its expression levels could affect cell morphology. As shown in Fig. 2A, without Dox, cells retained their classical morphology of PSC, growing as small round and compact cells in colonies with well-defined edges. However, silencing of E-cadherin caused a gradual change to a spindle-like morphology and produced a clear alteration in cellular distribution since each cell became isolated from other neighboring cells, as can also be seen in the nuclear density in Figure 1A. All these morphological alterations were not detected in parental HES3 incubated with or without Dox (Fig. S3). Since major morphological changes were observed in cells incubated with Dox at 96 h, we decided to measure cell area compared to control cells. Quantification of this parameter confirmed our qualitative observations. Values of cell area were higher in cells cultured with Dox compared to cells without Dox (control condition). Thus, these data allow us to confirm that E-cadherin downregulation produced morphological changes on HES3-KRAB^CDH1^ cells.

**Fig. 2.**
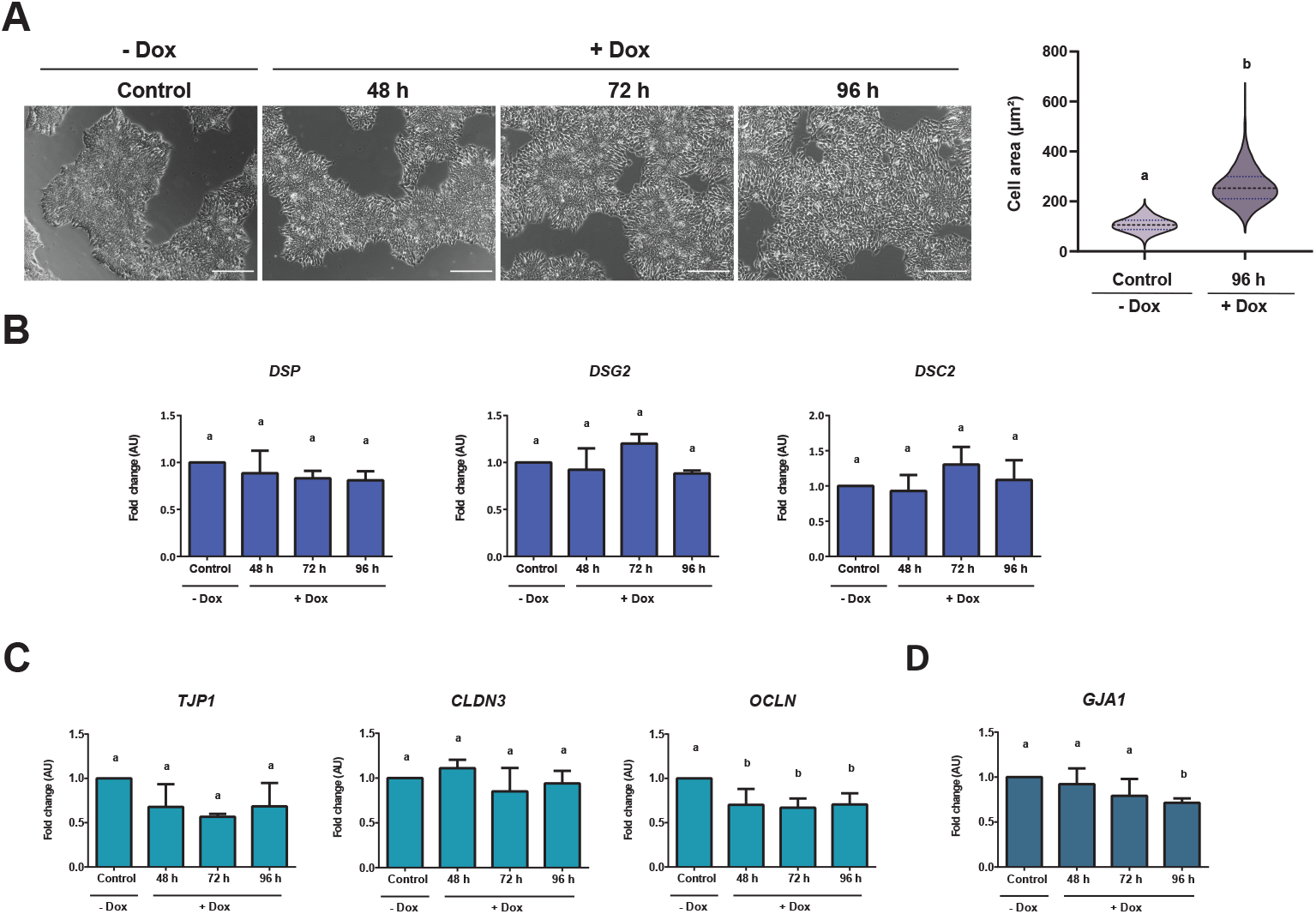
Downregulation of E-cadherin induced morphological alterations and transcriptional regulation of some adhesion complexes genes. A) Left panel: Representative images of HES3-KRAB^CDH1^ cells incubated with Dox at different times. Scale bar 200 μm. Right panel: quantification of cell area in HES3-KRABCDH1. Cells were incubated without Dox (Control) or with Dox during 96 h (96 h). Area was measured for each cell individually. B) Relative mRNA expression of desmoplakin (*DSP*), desmoglein (*DSG2*), desmocollin (*DSC2*) C) tight junction protein 1 (*TJP1*), claudin 3 (*CLDN3*), occludin 1 (*OCLN*) and D) connexin 43 (*GJA1*), when E-cadherin is downregulated. AU is the abbreviation for arbitrary units. Results are represented as mean ± SD (n=5). ANOVA with post hoc Tukey. Different letters indicate significant differences of groups compared to control condition (p<0,05).

First steps of EMT involve destabilization and disassembly of cell-cell contact complexes including desmosomes, adherens, tight and gap junctions. To explore the idea that morphological changes induced by E-cadherin downregulation could be related to disassembly mediated by transcriptional regulation of other junction structures, we evaluated mRNA levels of several genes that compose this complex. Reduction of E-cadherin did not induce any changes in the expression of genes involved in desmosome junctions such as desmoplakin (*DSP*), desmocollin (*DSC2*) and desmoglein (*DSG2*) (Fig. 2B). Also, no significant changes were observed in the expression of tight junction protein 1 (*TJP1*) and claudin 3 (*CLDN3*) two genes implicated in tight junctions (Fig. 2C). However, following E-cadherin downregulation, occludin 1 (*OCLN*), another gene implicated in this type of junction, showed a significant decrease. In the same line, expression of connexin-43 (*GJA1*), a primary component of gap junctions, also was significantly decreased compared to control condition (Fig. 2D). Together, data demonstrated that expression of some components of intercellular junctions are modulated, suggesting that E-cadherin silencing could be inducing a partial alteration of genes related to the maintenance of cell structure.

### E-cadherin silencing activate genes related to initiation of EMT

As our previous results suggested that E-cadherin downregulation induce some alterations that might be related to early stages of EMT, our aim was to evaluate if this decrease could initiate, activate and/or maintain this transition process. For this, we performed a characterization of the expression pattern of the pluripotency genes *OCT4* (*POU5F1*), *NANOG*, *LIN28* (*LIN28A*), genes involved in EMT namely *SNAI1*, *SNAI2*, *ZEB1* and *ZEB2*, and the genes of early mesoderm precursor *TBX6*, *MIXL1* after Dox incubation and consequent E-cadherin decrease. As shown in Fig. 3A, *OCT4* mRNA expression was significantly reduced only at 72h, while no changes in *NANOG* and *LIN28* expression were observed. Although gene expression of transcription factors ZEB1 and ZEB2, as well as early mesoderm genes did not present significant differences, we found that expression of SNAI1 and SNAI2, two key master transcription factors in the initiation of EMT were significantly upregulated (Fig. 3B and C).

**Fig. 3.**
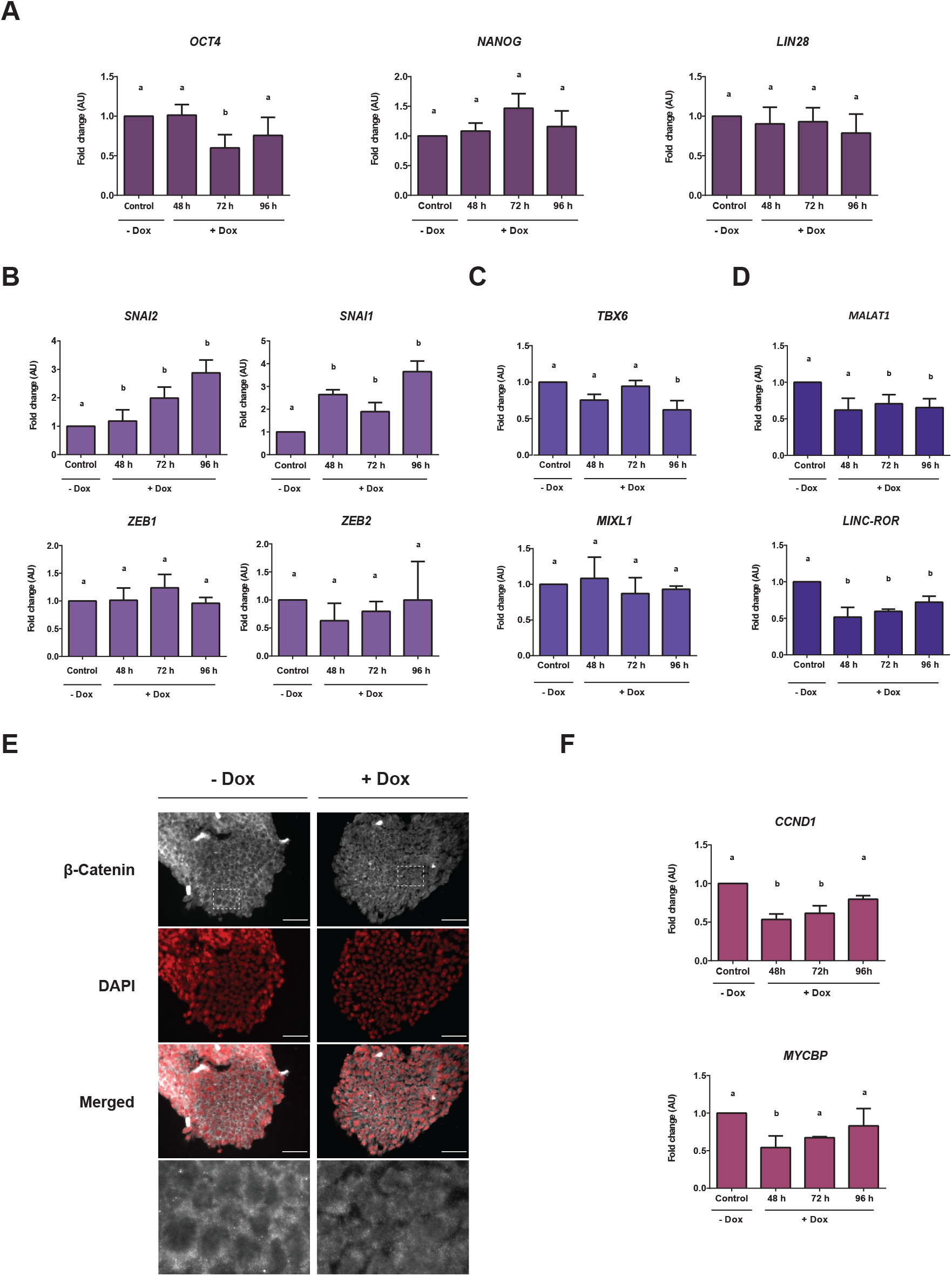
E-cadherin decrease activated the initiation of EMT. qPCR analysis of A) genes of pluripotency *OCT4*, *NANOG* and *LIN28*, B) genes involved in EMT such as *SNAI1*, *SNAI2*, *ZEB1* and *ZEB2* and C) genes of early mesoderm precursor *TBX6* and *MIXL1* when E-cadherin is downregulated. D) Relative expression level of *LINC-ROR* and *MALAT1*, two lncRNAs that act as regulators of EMT were measured by qPCR. AU is the abbreviation for arbitrary units. Results are represented as mean ± SD (n=5). ANOVA with post hoc Tukey. Different letters indicate significant differences of groups compared to control condition (p<0,05). E) Immunofluorescence of b–catenin (red) in cells treated with Dox. Nuclei were stained with DAPI. Magnification of the indicated areas are shown in black and white colors. Representative images of at least three experiments are shown. Scale bar 50 μm. F) Relative mRNA levels of c-myc (*MYCBP*) and cyclin D1 (*CCND1*) were determined by qPCR. Data are presented as mean ± SD (n=4). ANOVA with post hoc Tukey. Different letters indicate significant differences of groups compared to control condition (p<0,05).

Activation of EMT occurs in response to various factors and signals. Different reports tend to focus on the study of long non coding RNA (lncRNA) as fundamental regulators of EMT in cancer stem cells and PSC (16). Based on this background, our next step consisted on evaluating the expression of *LINC-ROR* and *MALAT1*, two lncRNAs considered as EMT regulators, during E-cadherin downregulation. As observed in Fig. 3D, both lncRNAs were significantly down-regulated in all experimental conditions. Another factor that contributes to EMT activation is β-catenin, a protein that interacts with the cytoplasmic domain of E-cadherin. When the latter is downregulated, β-catenin is released into the cytoplasm and can translocate into the nucleus (17), leading to transcriptional activation of genes related to EMT. In order to have an approximation about activation of this pathway, we evaluated localization of β-catenin after E-cadherin downregulation. Immunofluorescence showed that, in control conditions, β-catenin appears to be localized in the cell membrane. Upon reduction of E-cadherin, we observed an accumulation of this protein in the cytoplasm (Fig 3E). Since we only could observe a delocalization of this protein, to determine if β-catenin effectively translocated into the nucleus, we analyzed the expression of two target genes of β-catenin signaling pathway, cyclin D1 (*CCND1*) and c-myc (*MYCBP*) (Fig 3F). Expression levels of both genes decreased at 48 and 72 h of Dox treatment, but returned to basal levels at 96 h suggesting that β-catenin pathway was not activated after E-cadherin silencing. All together, these results indicate that even though other factors and signals that drives EMT do not seem to be activated, E-cadherin downregulation could be triggering the initial steps of EMT.

### Downregulation of E-cadherin induces a partial EMT state

Rather than a binary switch between two states, it is accepted that during EMT cells can transiently adopt different intermediate states collectively described as partial EMT. To test whether E-cadherin downregulation could be inducing a partial EMT state, some features of this particular state were analyzed. As previously observed, both 72 and 96 h of Dox treatment induced a pronounced decrease of E-cadherin. In view of this, we decided to evaluate functional effects of E-cadherin silencing regardless of incubation time. While transitioning, cells can attain a partial EMT phenotype that enables them to move collectively which is characteristic of this state. Thus, we next evaluated cell migratory capacity through a wound healing assay. For this, cells were treated with Dox and then a scratch was performed. Parental HES3 wild type cells did not display any migratory difference upon Dox treatment, indicating that Dox treatment per se does not modify the migratory phenotype of cells (Fig. S4). Importantly, when we performed this experiment in HES3-KRAB^CDH1^ cells, we observed that Dox treatment induced a significant increase in wound closure, suggesting that down-regulation of E-cadherin stimulated collective cell migration (Fig. 4A).

**Fig. 4.**
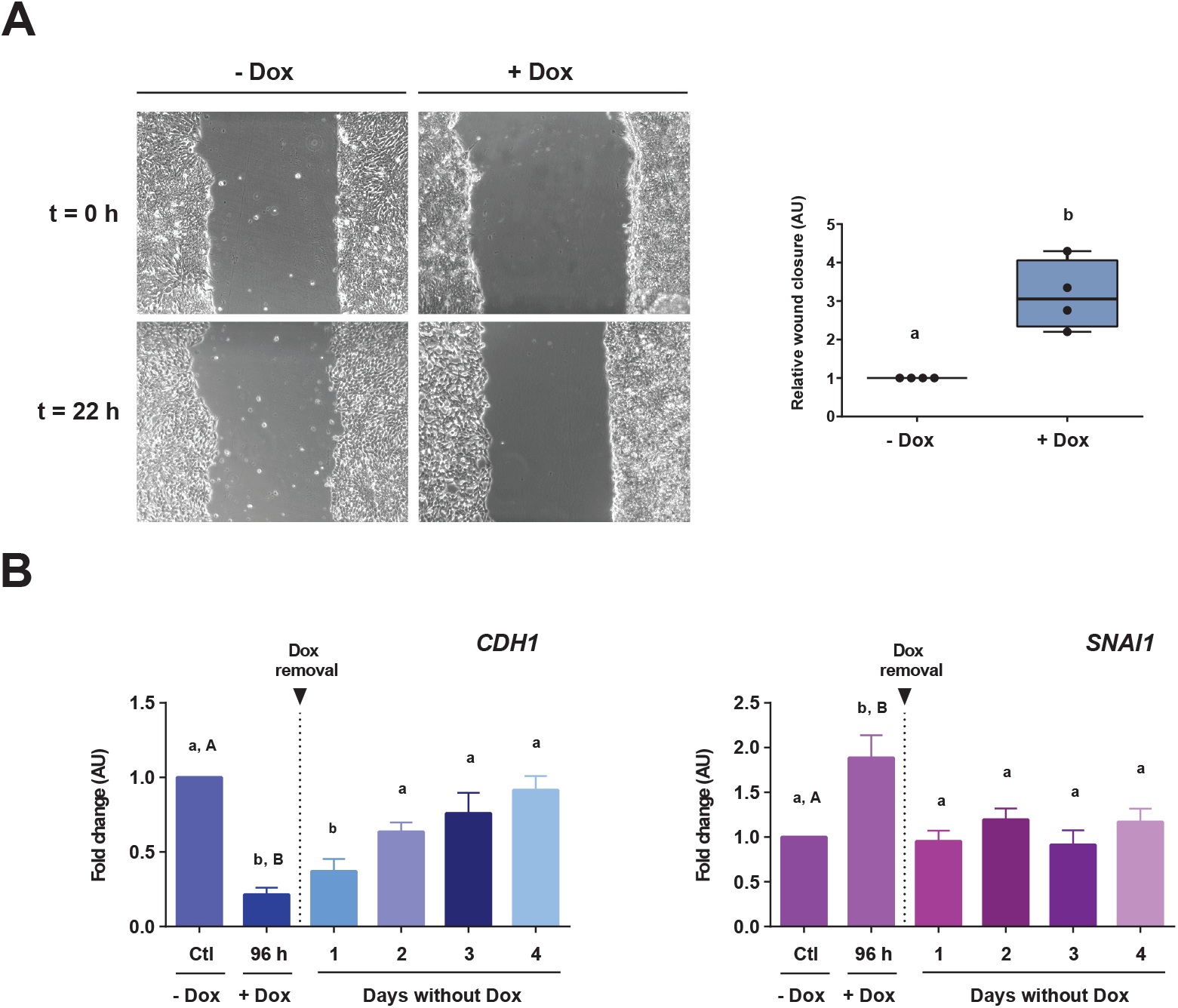
Silencing of E-cadherin induced a partial EMT state. A) Wound healing assay was made after cells were treated with or without Dox (+Dox and −Dox, respectively) (t=0h). Images were taken at t=0 h and after 22 h of recovery (t=22h) (left). Graphic represents the relative wound closure area of at least four independent experiments (right). AU is the abbreviation for arbitrary units. T-test. Different letters indicate significant differences. B) Rescue experiment to regain the knockdown effect. Cells were incubated with Dox for 96 h. At this time point, Dox was removed and cells were maintained for another 96h (4 days) only with culture medium. *CDH1* and *SNAI1* gene expression was evaluated by qPCR. Results are represented as mean ± SD (n=5). AU is the abbreviation for arbitrary units. ANOVA with post hoc Tukey. Different letters indicate significant differences of groups compared to control condition (p<0,01).

Hybrid phenotype of partial EMT is usually described as metastable and since acquisition of this state is transitory, cells can revert to a previous state leaving in evidence their high plasticity. As an approach to evaluate reversibility of partial EMT phenotype, we performed a rescue experiment where cells were treated with Dox for 96 h. Afterwards, Dox was removed and cells were maintained only with medium for other 96 h (4 days). When Dox was removed, both E-cadherin downregulation and increase in *SNAI1* expression were reversed (Fig. 4B). mRNA levels of E-cadherin returned slowly to the same levels of control conditions within two days of Dox removal. Moreover, *SNAI1* expression returns to control levels the following day after Dox removal. These findings indicate that the effect of E-cadherin silencing is reversible, showing flexibility and plasticity of cells in partial EMT.

## Discussion

EMT is a complex mechanism that involves morphological and molecular changes by which epithelial cells can acquire a mesenchymal phenotype. In this work we investigated the molecular effects of E-cadherin (CDH1) downregulation as a triggering signal for EMT activation in hESC. Overall, we found that this alteration induces a partial EMT where cells attain a hybrid state retaining their undifferentiated state while showing certain features of a mesenchymal-like phenotype.

In hESC, cell differentiation involves the loss of the undifferentiated state, activation of EMT and acquisition of a specific lineage. Molecular characterization of HES3-KRAB^CDH1^ following E-cadherin knockdown showed a transient disturbance in *OCT4* expression. A precise level of this gene must be sustained for the maintenance of pluripotency (18) and small variations could be related to this balance. Since no changes in *NANOG* and *LIN28* expression were observed, our data suggest that pluripotency of these cells was not disturbed. However, E-cadherin silencing did induce a marked increase in *SNAI1* and *SNAI2* levels, alterations that were not observed in previous reports where E-cadherin abrogation in embryonic stem cells was evaluated (19). Experimental observations reported that SNAI1 is induced at the initiation of EMT whereas ZEB1 is produced at a lather phase. In fact, SNAI1 is proposed to trigger this process inducing a decrease in E-cadherin followed by a second step where ZEB1 further reinforces this effect (20–23). These sequential steps are necessary for a complete EMT to occur. Induction of *SNAI1*, but not of *ZEB1* as observed in our results, suggest that E-cadherin partial downregulation does not have enough strength on its own to signal the next step of EMT resulting in an incomplete event. This could be explained by the regulation of SNAI1 and E-cadherin expression. E-cadherin prevents stimulation of NF-appa-B and other signaling pathways, but when its levels are low this pathway is activated and acts as an inductor of *SNAI1* expression (24). Moreover, SNAI1 levels are amplified by a self-stimulatory loop resulting in higher levels of this gene which are necessary for activation of other transcriptional repressors of E-cadherin such as ZEB1 (22). Probably, in our system, partial E-cadherin downregulation did not induce an increase of *SNAI1* to the levels necessary to activate *ZEB1*, resulting in a partial EMT. Currently there is no general agreement on the molecular markers that identify univocally cells in a partial EMT state. Currently, some reports have determined partial EMT state based on the detection of E-cadherin and vimentin (25), markers of epithelial and mesenchymal state, respectively.

On the other hand, Puram *et al* (26) analyzed a gene expression panel for features of EMT detecting an increased expression of TGF-b (a regulator of EMT) and vimentin while the expression of epithelial markers and other classical EMT transcription factors remain unaltered. In view of these findings, we considered a wide range of molecules and features of EMT to make a complete molecular characterization. Our results indicated that E-cadherin silencing in HES3-KRAB^CDH1^ cells caused a higher expression of EMT transcription factors genes *SNAI1* and *SNAI2*, an alteration of cell morphology to a mesenchymal-like phenotype and an enhanced collective cell migration. Notably, all these partial changes correspond to those observed during EMT. However, several other reports have shown that SNAI1 functions are not restricted only to EMT initiation. This transcription factor participates in all the events mentioned above, inducing morphological changes, regulating cell adhesion and partially modulating genes of tight junctions (27, 28). This suggests that SNAI1 also could be implicated in these alterations and in the acquisition of the incomplete EMT state.

Even though loss of E-cadherin is considered a hallmark event of EMT, many other signals and cellular events are required to activate this process. Up to recent years, lncRNAs has emerged as important regulators of several biological processes (16). *LINC-ROR* is highly expressed in ESC and its implicated in the maintenance of pluripotency since act as sponge of microRNA that repress the translation of pluripotency genes, ensuring the undifferentiated state. When *LINC-ROR* expression decreases, synthesis of these pluripotent genes is repressed. However, a positive feedback loop of this pluripotency genes enables them to control and activate their own synthesis (29). This autoregulatory circuitry provides a mechanism by which stem cells retain their ability to react appropriately to differentiation signals (30). Our data suggest that although E-cadherin decrease down regulated *LINC-ROR* levels, it does not have enough strength as individual signal to induce a disturbance in pluripotency genes expression. On the other hand, β-catenin signaling is determined by its stabilization and accumulation in the cytoplasm (17). In fact, a key step in the WNT pathway is regulation of β-catenin cytosolic pool which is available for nuclear translocation and activation of gene transcription (31).

A partial downregulation of E-cadherin as occurs in our system could involve lower free β-catenin levels than threshold levels necessary to translocate into the nucleus and activate this pathway.

As concluding remarks, recent experimental findings have bolstered the relevance of partial EMT until to be considered as a focal point of study in the EMT field. Due to their unstable nature, it has been extremely difficult to establish adequate models that enable to have a better understanding about this particular state. In this work, we could establish a model of partial EMT in hESC induced by downregulation of E-cadherin, a hallmark event of EMT. Moreover, this is the first time that this intermediary state is detected in PSC. We believe our results can be used as a platform to identify different aspects of this particular state. Results obtained here provide a useful tool that will allow us to investigate molecular events of an unstable intermediate state and identify multiple layers of molecular changes that occur during partial EMT.

## Acknowledgements

This work was supported by grants from the National Agency for Scientific and Technical Promotion (ANPCyT) and from the Scientific and Technical Research Fund (FONCyT) PICT-2015-0868. Authors would like to thank FLENI-CONICET and Pérez Companc Foundation for their continuous support.

## Competing Interests

The authors declare that they have no competing interests.

## Supplementary Material

**Table S1.**
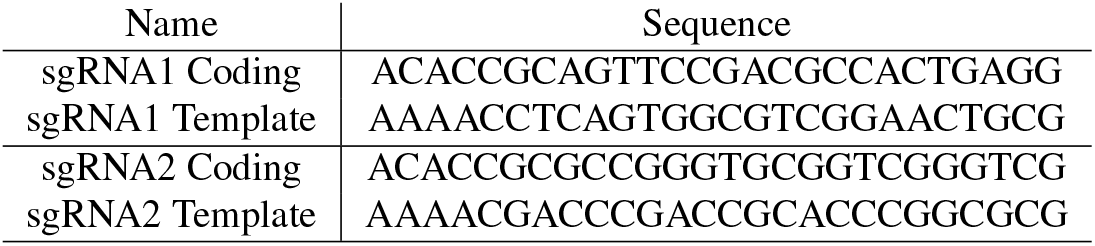
sgRNA sequences

**Table S2.**
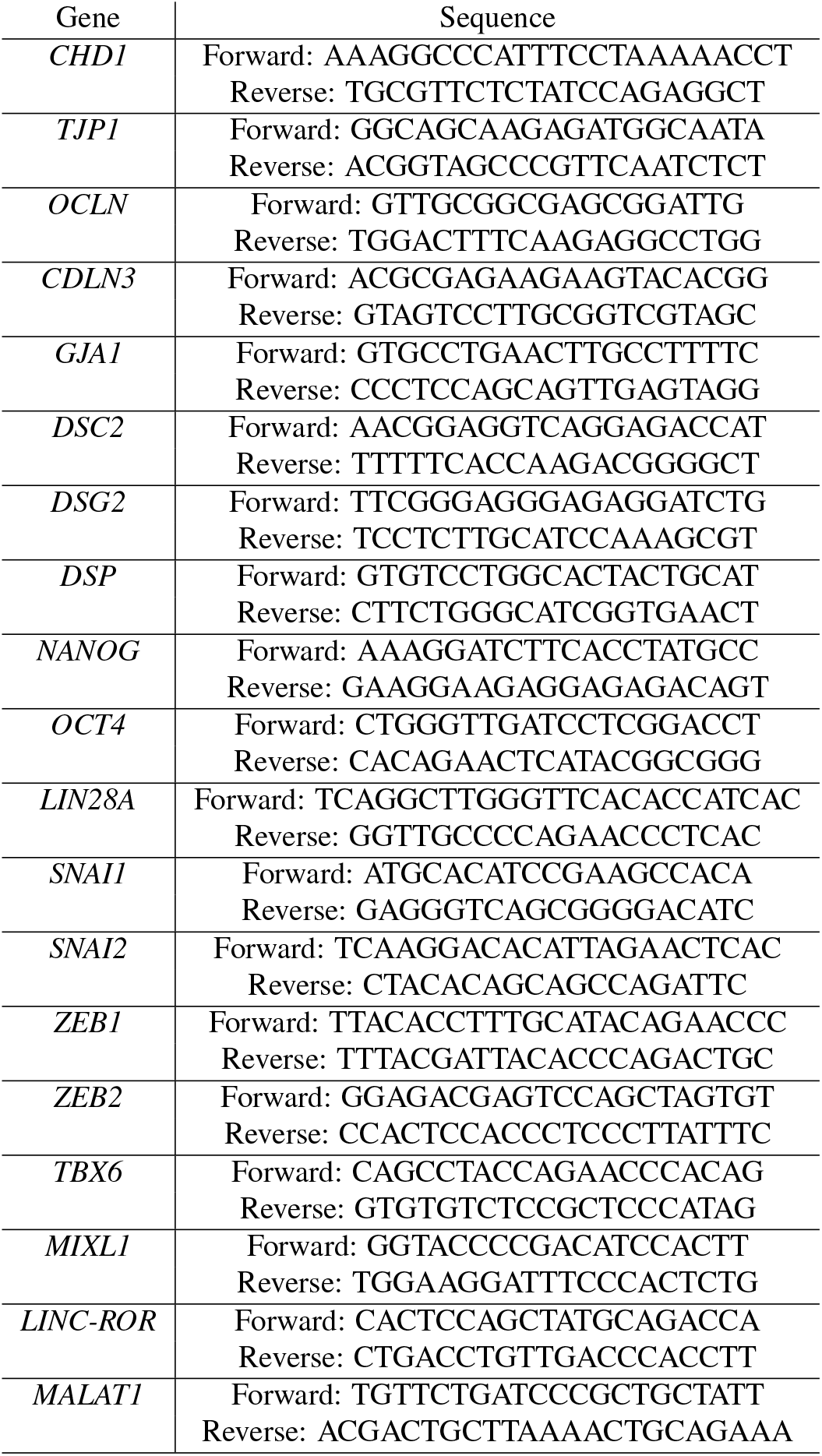
Primers sequences

**Fig. S1.**
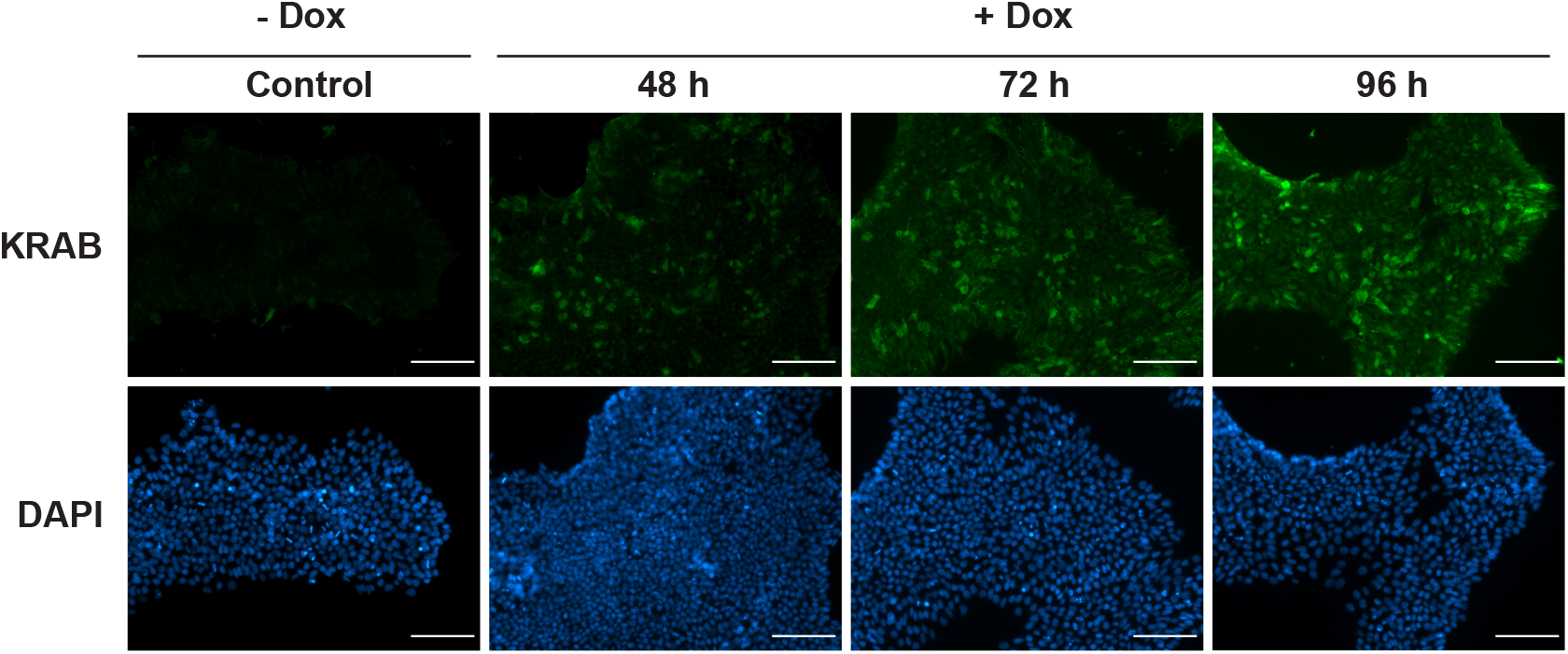
Expression of KRAB increased gradually after Dox incubation. Immunofluorescence images of KRAB protein expression in cells incubated with Dox during 48, 72 and 96 h. Nuclei were stained with DAPI. Representative images of at least three experiments are shown. Scale bar 100 μm.

**Fig. S2.**
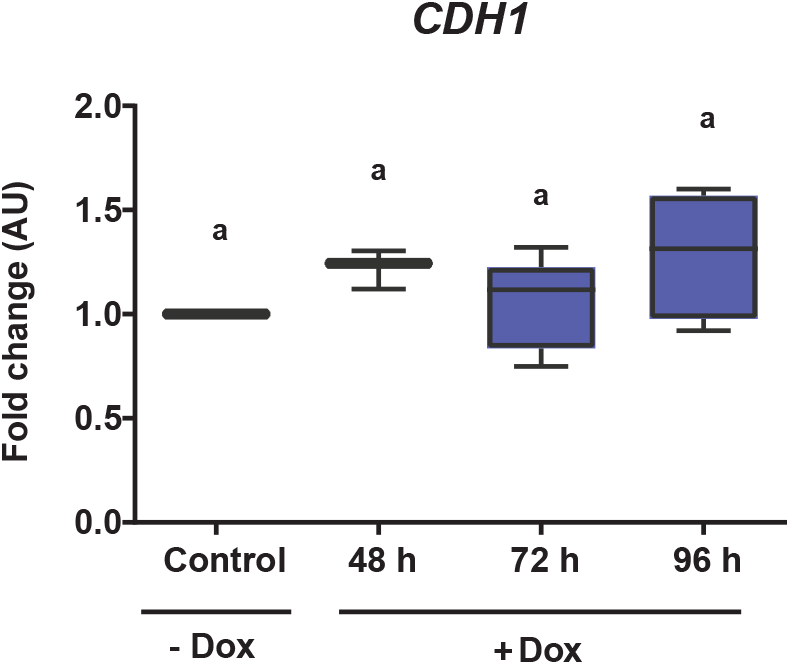
E-cadherin levels were not affected in parental HES3 cells after Dox incubation. Relative mRNA levels of CDH1 in parental HES3 wild type cells after incubation with Dox at different times assessed by qPCR. AU is the abbreviation for arbitrary units. Results are represented as mean ± SD (n=5). Different letters indicate significant differences of groups compared to control condition for ANOVA with post hoc Tukey.

**Fig. S3.**
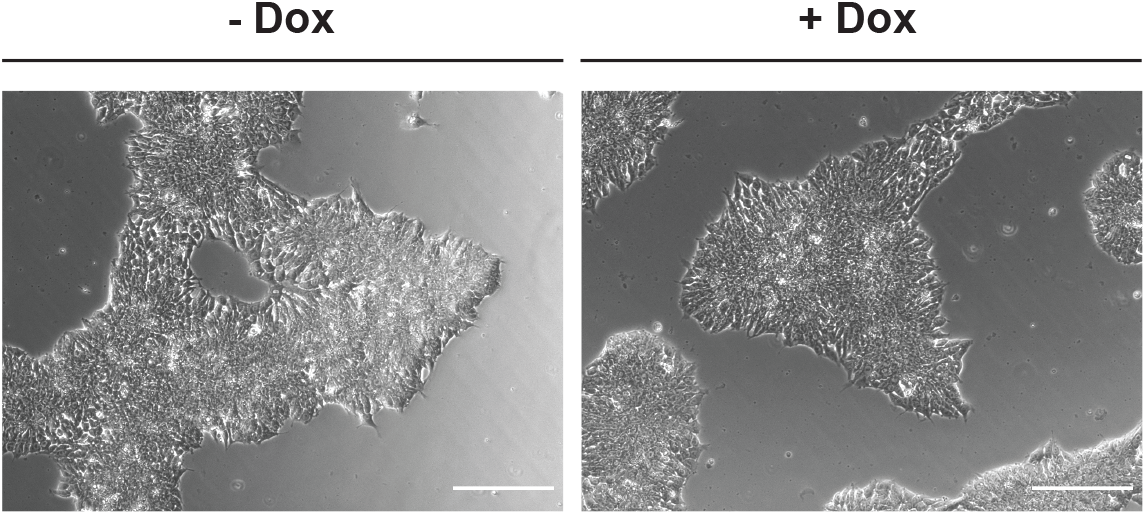
Cell morphology was not affected in parental HES3 cells incubated with Dox. Representative images of parental HES3 wild type cells incubated with Dox at different times. Scale bar 200 μm.

**Fig. S4.**
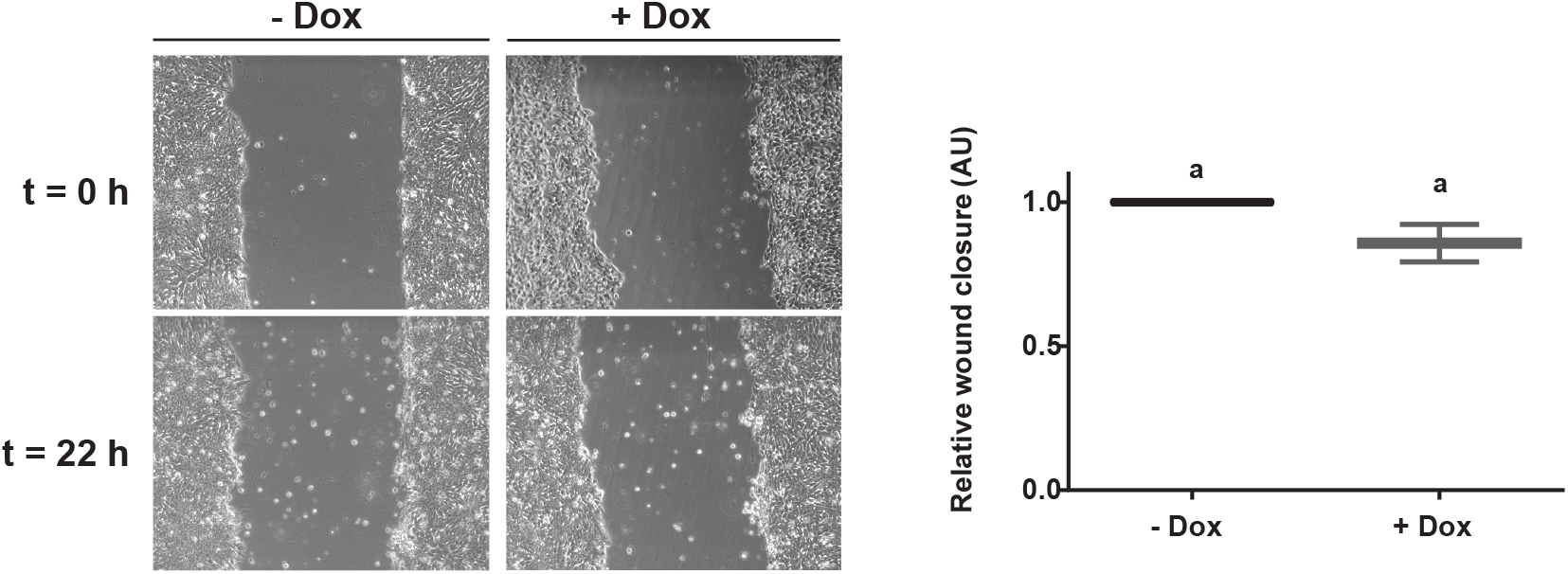
Cell migration was not enhanced in parental HES3 cells after Dox incubation. Wound healing assay was made after parental HES3 wild type cells were treated with or without Dox (+Dox and −Dox, respectively) (t=0h). Representative images of at least three experiments are shown. Images were taken at t=0 h and after 22 h of recovery (t=22h).

## Notes

### Competing Interest Statement

The authors have declared no competing interest.

### Summary of Updates

Figure 2A updated. Cell area quantification was re-analyzed using Cellpose tool to make automatic masks for posterior ImageJ-FIJI analysis. Corresponding method and citation was added as well.

